# Strigo-D2 – a bio-sensor for monitoring the spatio-temporal pattern of strigolactone signaling in intact plants

**DOI:** 10.1101/2021.08.03.454859

**Authors:** Changzheng Song, Jiao Zhao, Marjorie Guichard, Dongbo Shi, Guido Grossmann, Christian Schmitt, Virginie Jouannet, Thomas Greb

## Abstract

Strigolactones (SLs) are a class of plant hormones modulating developmental programs in response to endogenous and exogenous stimuli and mediating biotic interactions. However, a comprehensive view on the spatio-temporal pattern of SL signaling has not been established and tools for a systematic *in planta* analysis do not exist. Here, we present Strigo-D2, a genetically encoded ratiometric SL signaling sensor, allowing the examination of SL signaling distribution with cellular resolution and its rapid response to altered SL levels in intact plants. By monitoring the abundance of a truncated and fluorescently labeled SUPPRESSOR OF MAX2 1-LIKE 6 (SMXL6) protein, a proteolytic target of the SL signaling machinery, we show that all cell types investigated have the capacity to respond to changes in SL levels but with very different dynamics. In particular, SL signaling is pronounced in vascular cells but low in guard cells and the meristematic region of the root. We also show that other hormones leave Strigo-D2 activity unchanged indicating that initial SL signaling steps work in isolation from other hormonal signaling pathways. Specificity and spatio-temporal resolution of Strigo-D2 underline the value of the sensor for monitoring SL signaling in a broad range of biological contexts and with highly instructive analytical depth.

## Introduction

Strigolactones (SLs), a class of carotenoid-derived phytohormones, were originally identified in plant root exudates acting as germination stimulants for parasitic plants [1]. Since then, an increasing number of roles of SLs as stimulants of biotic interactions [2] and as endogenous growth regulators in a broad range of species has been unveiled. The spectrum of SL-dependent processes includes the determination of root architecture, shoot branching, radial growth, leaf development, flower size, and the adaptation to drought and nutrient availability [3-6] identifying SL biology as a highly relevant topic for exploring and modifying plant performance. In spite of this broad spectrum of SL-dependent processes and the role of SLs in long-distance communication [7], knowledge about SL distribution and the pattern of SL signaling with high spatial resolution is surprisingly scarce.

Biosynthesis of SLs is initiated by converting all-trans-β-carotene to 9-cis-β-carotene through the all-trans/9-cis-carotene isomerase DWARF27 (D27). The 9-cis-β-carotene is then sequentially converted by CAROTENOID CLEAVAGE DIOXYGENASE 7 (CCD7) and CCD8 homologs into carlactone which is the last common biosynthetic precursor for all known SLs [8, 9]. The downstream biosynthetic pathways leading to functional SLs vary among species, resulting into species-specific SL repertoires [10-12]. In all cases, however, canonical SLs carry a butenolide ring (D-ring) linked to a less conserved tricyclic lactone (the ABC rings) via an enol-ether bond [13, 14]. Based on stereochemical differences at the junction of the B and C rings, SLs are divided into strigol- and orobanchol-like subfamilies, exemplified by 5-deoxystrigol and 4-deoxyorobanchol, respectively. In addition to canonical SLs, there are compounds with SL-like activity lacking the B-and C-rings such as methyl carlactonoate (MeCLA) and heliolactone [13, 14]. Overall, at least 25 different compounds with SL-like activity have been discovered in plants all exhibiting a 2’R configuration in the D-ring [14]. Importantly, grafting experiments and gene expression analyses indicate that SLs are synthesized in both roots and shoots [15-17] but can be transported long distances possibly along the vasculature [18].

In Arabidopsis, similarly as in other species, bioactive SL molecules are perceived by homologs of the nuclear α/β-hydrolase superfamily protein DWARF14 (D14) [19-21]. Although still being controversial in some details, the current view is that D14 proteins fulfill a role as both SL receptors and as enzymes cleaving and deactivating SL molecules [21, 22]. In any case, binding of SLs to D14 induces conformational changes in the D14 structure and its recruitment to an Skp, Cullin, F-box (SCF) E3 ubiquitin-protein ligase complex, containing the F-box protein MORE AXILLARY GROWTH 2/DWARF3 (MAX2/D3) [21-23]. MAX2/D3 serves as an adapter providing specificity toward the recruitment of D14 and, upon D14 binding, also recruiting proteins from the SUPPRESSOR OF MAX2 1-LIKE/DWARF53 (SMXL/D53) family, in particular SMXL6, SMXL7, and SMXL8 [24, 25]. After final formation of an SCF^MAX2/D14/SMXL^ complex, SMXL proteins are polyubiquitinated and degraded by the cellular proteasome – the decisive step for modulating SL-dependent processes [26, 27].

*MAX2* is broadly expressed in the seedling stage, but predominantly in vascular tissues and meristems at adult stages [28-30]. *D14* expression patterns largely overlap with those of *MAX2* [30, 31] allowing physical interactions between these proteins and SL signaling in the nuclei of respective cells. Therefore, based on expression analyses of key signaling components, differences in the potential of different cell types to perceive and to transmit SL signals is likely. However, differences in signaling capacities among cell types and growth stages have not been investigated systematically so far.

Of note, the Arabidopsis KARRIKIN-INSENSITIVE2 (KAI2) protein, a close homolog of D14, mediates signal transduction of smoke-derived karrikin (KAR) molecules via a comparable SCF complex-based mechanism and, most likely, serves as a receptor for a yet to be identified endogenous signaling molecule [12, 20, 24, 32, 33]. As for D14 during SL signaling, MAX2 is responsible for recruiting KAI2 after KAR binding. Instead of SMXL6, SMXL7 and SMXL8, however, predominantly the SMXL family proteins SUPPRESSOR OF MAX2 1 (SMAX1) and SMXL2 are targeted by KAR signaling [24, 34, 35]. With regard to the specificity of the SL and KAR signaling pathways, it is important to be aware that synthetic SL analogs do not only hold a 2’R configuration but also an enantiomeric 2’S configuration and that the latter have the potential to activate KAI2 [36, 37]. Therefore, sharing components between SL and KAR signaling pathways and a lack of specificity of some synthetic hormone analogs makes it sometimes challenging to determine the effect of each pathway individually.

As key mediators of SL responses, nuclear SMXL/D53 proteins are predicted to contain an N-terminal double Clp-N domain (D1) and one or two P-Loop NTPase domains (D2, Fig 1A) [22, 38]. Although their exact mode of action in the nucleus is unclear, SMXL6, SMXL7 and SMXL8 proteins show DNA binding capacities *in vitro* and *in planta* and determine the activity of decisive regulatory genes like, for example, *BRANCHED1* (*BRC1*) in the context of shoot branching [39]. Importantly, previous research by Shabek and colleagues [22] showed that the D2 domain of the rice D53 protein alone forms a stable complex with D14-D3-ASK1 proteins and is also degraded by the proteasome in a MAX2/D3-dependent manner. This indicates that the D2 domain is sufficient for hormone-induced and SCF^MAX2/D14^-catalysed protein turnover.

**Fig 1.**
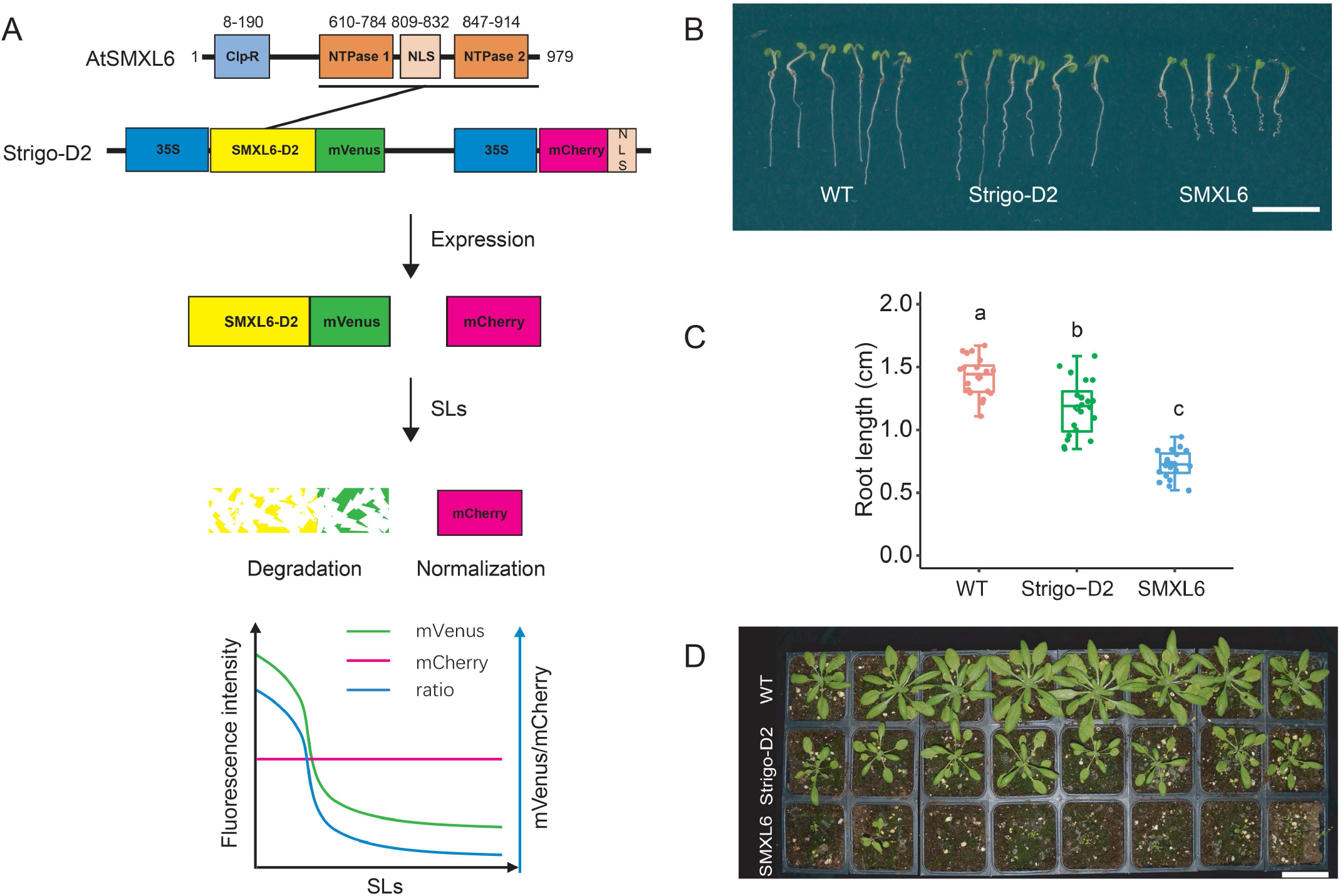
Sensor construction and phenotypic analysis of Strigo-D2 lines. **(A)** Schematic representation of the SMXL6 protein domain structure and the Strigo-D2 design. Numbers above domains represent amino acid positions. The D2 domain ranges from 615 to 979 aa. **(B-D)** Phenotypes of transgenic plants expressing SMXL6-D2-mVenus (shown is line 1 mentioned in Fig. 2) and SMXL6-mVenus in comparison to wild type plants (WT). (B) Seedlings five days after germination. (C) Root length quantification of five day-old seedlings. Different letters above each box indicate statistical groups (One-way ANOVA, *p* < 0.0001, n = 20). (D) Plants four weeks after germination. Scale bars: 1 cm (B) or 5 cm (D).

Genetically encoded fluorescent biosensors are versatile tools for detecting changing levels of small molecules *in vivo* [40, 41]. With the unexcelled advantage of revealing relative molecule levels with high spatio-temporal resolution and within a physiological range, sensors have been developed for measuring small molecules like calcium, sugars and various plant hormones [42-46]. For SLs, ratiometric biosensors exploiting the levels of the SMXL6 or SMXL7 proteins as indication for the activity of the SL signaling pathway were characterized previously [35, 47-49]. The sensors responded as expected to various natural or synthetic SLs with high specificity, sensitivity and quantitative resolution demonstrating the suitablity of using proteolytic targets of the SL signaling pathway to estimate respective signaling levels. However, described sensors use partly bioluminescence as a readout hampering cellular imaging and requiring continuous supply of luciferin, the substrate of the luciferase reporter [35, 47, 48]. Moreover, ratiometric SL sensors have only been tested successfully in transient expression systems so far [35, 47-49] leaving open the question towards their performance in stable transgenic plant lines and in the context of an SL-controlled process. In comparison, a sensor stably integrated in plants expressing the luminescently labeled D2 domain of SMAX1 under the control of the ubiquitously active *UBQ10* promoter was shown recently to faithfully report KAR signaling (*pRATIO2251-SMAX1*_*D2*_ [35]). Demonstrating the challenges of these apporaches, the fluorescent reference protein supposedly co-translated with the same sensor could not be detected, though, preventing normalization of sensor activity [35].

Here, we present the fluorescent ratiometric biosensor Strigo-D2 allowing semi-quantitative monitoring of SL signaling levels with cellular resolution in intact Arabidopsis plants (Fig 1A). Strigo-D2 employs ubiquitous expression of the SMXL6 D2 domain fused to the yellow fluorescent protein mVenus to reveal the capacity of cells to proteolytically degrade SL signaling targets. SMXL6-D2-mVenus levels are directly compared to levels of the red fluorescent protein mCherry expressed under the control of the same ubiquitously active promoter from the same transgene allowing convenient normalization of signal intensities.

## Results

### Strigo-D2-plants show only mild growth alterations

To test the capacity of the SMXL6 protein to serve as an *in planta*-SL signaling sensor, we generated plant lines expressing full length SMXL6 or only the SMXL6-D2 domain fused to the yellow fluorescent mVenus protein [50] under the control of the broadly active *35S* promoter [51]. In addition, the lines expressed a *35S*-driven nuclear-localized version of mCherry [52] (mCherry-NLS) from the same transgene to normalize signal intensities (Fig 1A). Comparing transgenic plants with wild type revealed that plants expressing the full SMXL6-mVenus fusion protein showed much shorter roots at the seedling stage and substantially retarded growth as adults (Fig 1B-D), suggesting a severe interference of the broadly expressed SMXL6 protein with endogenous regulatory processes. Instead, roots of SMXL6-D2 expressing plants were only slightly shorter than wild type roots and growth was hardly affected overall (Fig 1B-D). When comparing fluorescence intensities, mCherry but no mVenus-derived signals were detected in nuclei of *p35S:SMXL6-mVenus_p35S:mCherry-NLS* seedlings expressing the full length SMXL6-mVenus protein using standard microscopy settings (Fig S1). In comparison, *p35S:SMXL6-D2-mVenus_p35S:mCherry-NLS* seedlings expressing SMXL6-D2-mVenus showed a clear nuclear mVenus signal when applying low laser intensity (Fig S1).

### Consistent (+)-5DS response in independent Strigo-D2 lines

Due to the mild effect of SMXL6-D2 on plant growth in comparison to the full-length protein and the robust detection of SMXL6-D2-mVenus fluorescence, we decided to explore whether SMXL6-D2-expressing plants are able to faithfully report on internal SL signaling levels. To this end, three independent transgenic SMXL6-D2 lines were treated with 0.5 μM (+)-5-deoxystrigol ((+)-5DS) inducing SL signaling [47, 53, 54]. In nuclei of the root maturation zone, a substantial reduction of the mVenus/mCherry intensity ratio was observed 10 minutes after application which continued to decrease for at least one hour (Fig 2A-2F). This indicated, as expected, a negative correlation between the mVenus/mCherry intensity ratio and the level of SL signaling. Changes in intensity ratios were solely dependent on a reduction of SMXL6-D2-mVenus levels as mCherry-derived signals were fully stable over time. These results demonstrated that the SMXL6-D2-mVenus protein responded to a pharmacological activation of SL signaling with a similar dynamics as previously reported for fluorescently labeled full length SMXL/D53 proteins [24, 27, 55]. Moreover, mCherry-NLS signals appeared to be a suitable reference for determining SL-dependent alterations in SMXL6-D2-mVenus levels. When comparing SMXL6-D2-mVenus signal dynamics among the three lines, line 1 showed the largest dynamic range over the incubation period (Fig 2A-F) prompting us to perform subsequent analyses taking advantage of the transgene present in this line. Expecting that the SMXL6-D2-mVenus/mCherry intensity ratio is tightly correlated with SL signaling levels, we named the sensor ‘Strigo-D2’ from here on.

**Fig 2.**
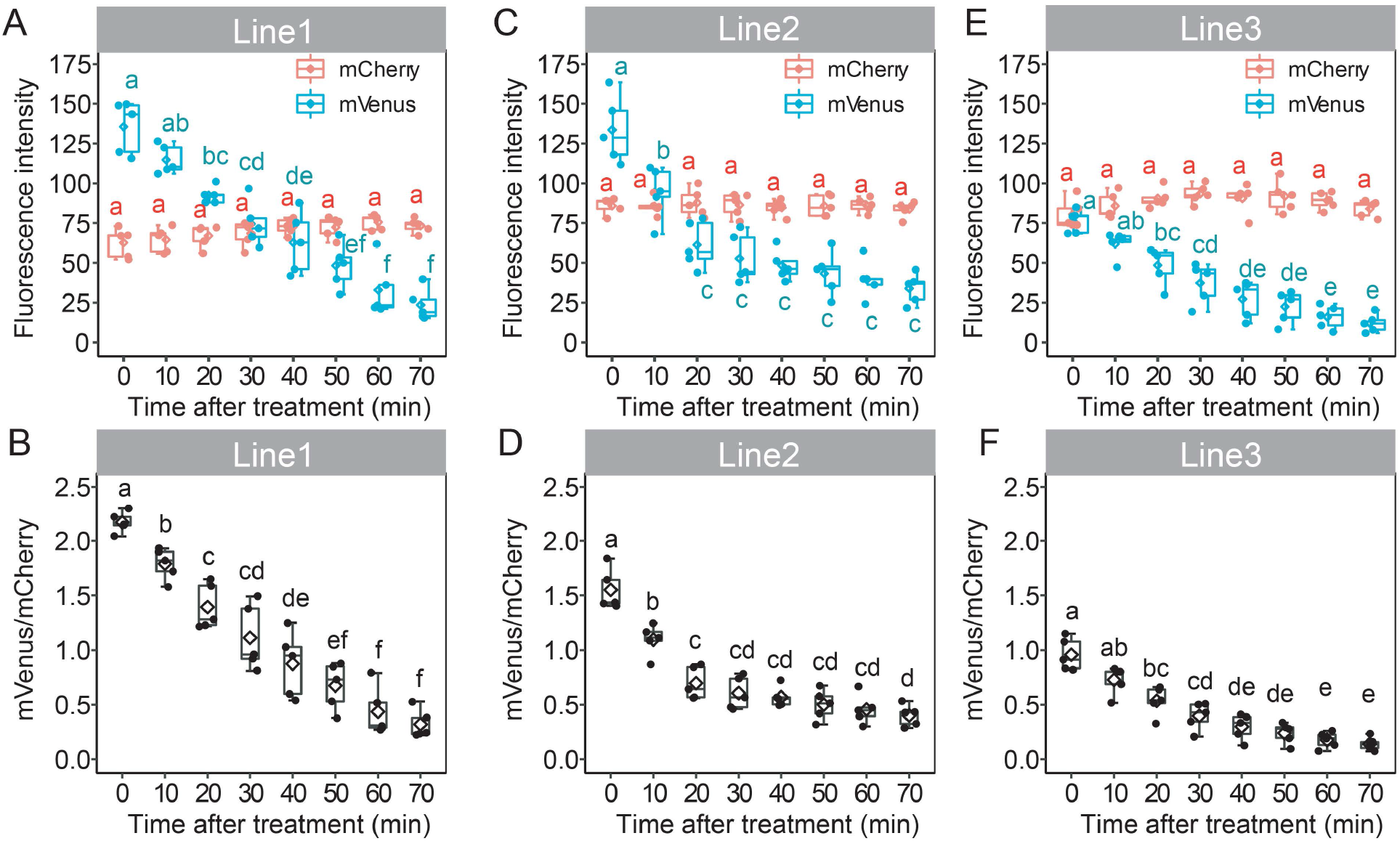
Strigo-D2 response to (+)-5DS in three independent transgenic lines. **(A-F)** Fluorescence intensity of mVenus and mCherry (A, C, E) and mVenus/mCherry signal ratios (B, D, F) in the root maturation zone upon treatment with 0.5 μM of (+)-5DS are shown for line 1 (A, B), line 2 (C, D), and line 3 (E, F). Different letters above error bars indicate statistical groups (One-way ANOVA, *p* < 0.01, n = 5). Each dot represents the value of one biological replicate as described in the methods section.

### The Strigo-D2 response depends on the SL signaling pathway

To see whether Strigo-D2 specifically reports on SL signaling, we treated Strigo-D2 plants for two hours with 0.5 µM (+)-5DS, *rac*-GR24, GR24^4DO^, or KAR1 [54]. As a result, a clear response was observed when applying the SL signaling inducers (+)-5DS, *rac*-GR24, and GR24^4DO^ but not when applying KAR1 which specifically induces KAR signaling [53, 54] (Fig 3A). In comparison, treating *d14* mutants carrying the same transgene had no effect on Strigo-D2 activity whereas the sensor responded like in wild type in *kai2* mutant backgrounds (Fig 3A). These results let us conclude that Strigo-D2 specifically responded to SL signaling and that this effect was fully dependent on the SL receptor D14. In this context it is important to note that *rac*-GR24 and (+)-5DS activate both strigolactone and karrikin signaling pathways whereas GR24^4DO^ specifically induces SL signaling [53, 54]. We could confirm this distinction by treating an *pSMXL5:SMAX1-Venus* reporter line [55] revealing KAR signaling responses (Fig 3A and Fig S2). Further supporting specificity of Strigo-D2 in reporting SL signaling levels, the phytohormones IAA, trans-Zeatin, ABA, and GA_3_, had no effect on sensor activity after two hours of exposure (Fig 3B and Fig S3). In summary, Strigo-D2 showed an SL-specific response, proposing it reliable for analyzing SL signaling at cellular resolution.

**Fig 3.**
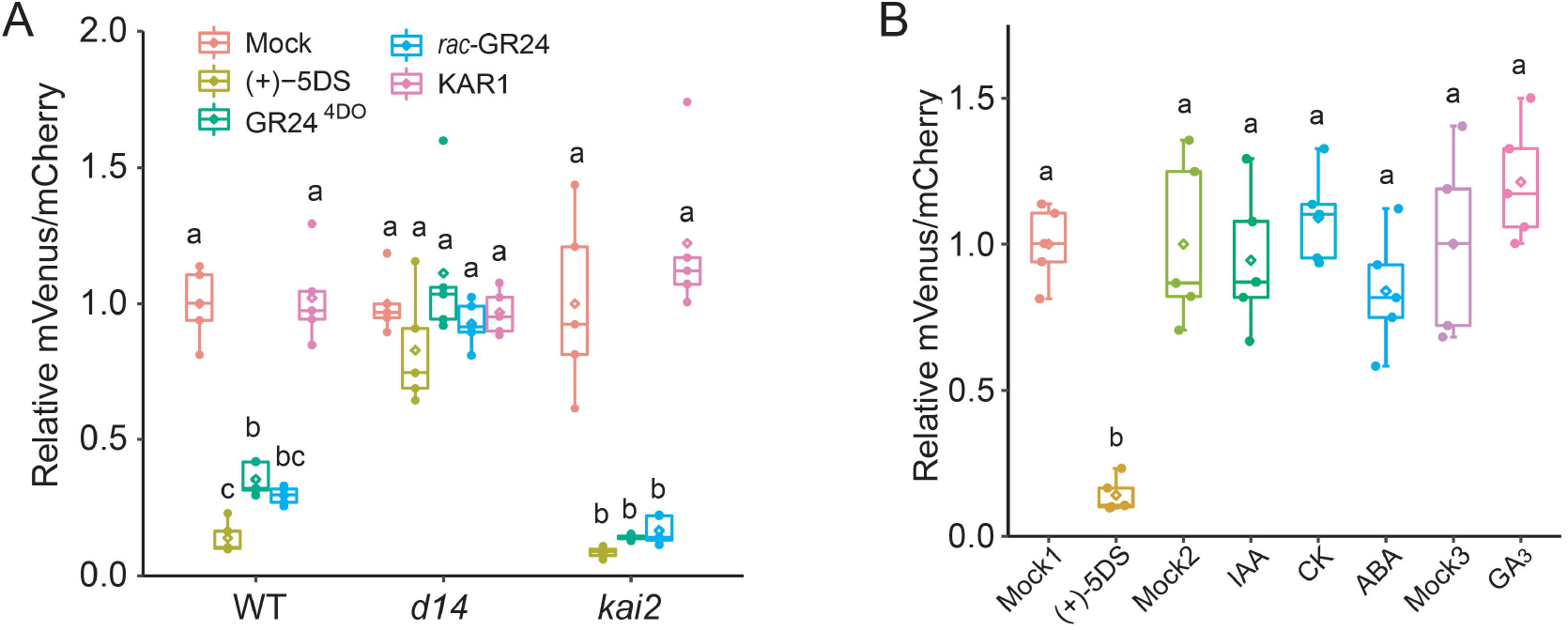
Specificity of Strigo-D2 in responding to the SLs signaling pathway. **(A)** Responses of Strigo-D2 in the root maturation zone of different genetic backgrounds (wild type, *d14, kai2*) to various SL molecules and to KAR1. **(B)** Responses of Strigo-D2 in the root maturation zone to other plant hormones (IAA, trans-Zeatin, ABA or GA3). Mock1: treatment with (+)-5DS solvent. Mock2: treatment with the solvent for IAA, trans-Zeatin, and ABA solvent. Mock3: treatment with the solvent for GA3. Letters above error bars indicate statistical groups (One-way ANOVA, *p* < 0.01, n = 5).

### Strigo-D2 shows cell type-specific activity patterns

To explore this potential, tile scanning of whole Strigo-D2 seedlings was performed revealing the activity pattern of Strigo-D2. As expected, signals of mVenus and mCherry were detected throughout 4-day-old seedlings, reflecting the ubiquitous expression of Strigo-D2 under the control of the *35S* promoter (Fig 4A-E, Fig S4). Interestingly, different intensity ratios of mVenus and mCherry fluorescence among tissues were found with ratios in meristematic and elongation zones of root tips being higher than those in root maturation zones and hypocotyls (Fig 4F-J, Fig S4). Intensity ratios were especially high in stomata guard cells which differed substantially from other epidermal cells in this regard (Fig 4F, J). In contrast, vascular cells displayed a low intensity ratio in all organs tested (Fig 4G, J, Fig S5) being in line with the reported vascular-associated expression of SL signaling components [29, 31] and as confirmed by fluorescent *MAX2* and *D14* promoter reporters (Fig S6). Taken together, we concluded that the activity of Strigo-D2 showed cell type-specific signatures and a spatial association of its activity with the expression pattern of SL signaling components.

**Fig 4.**
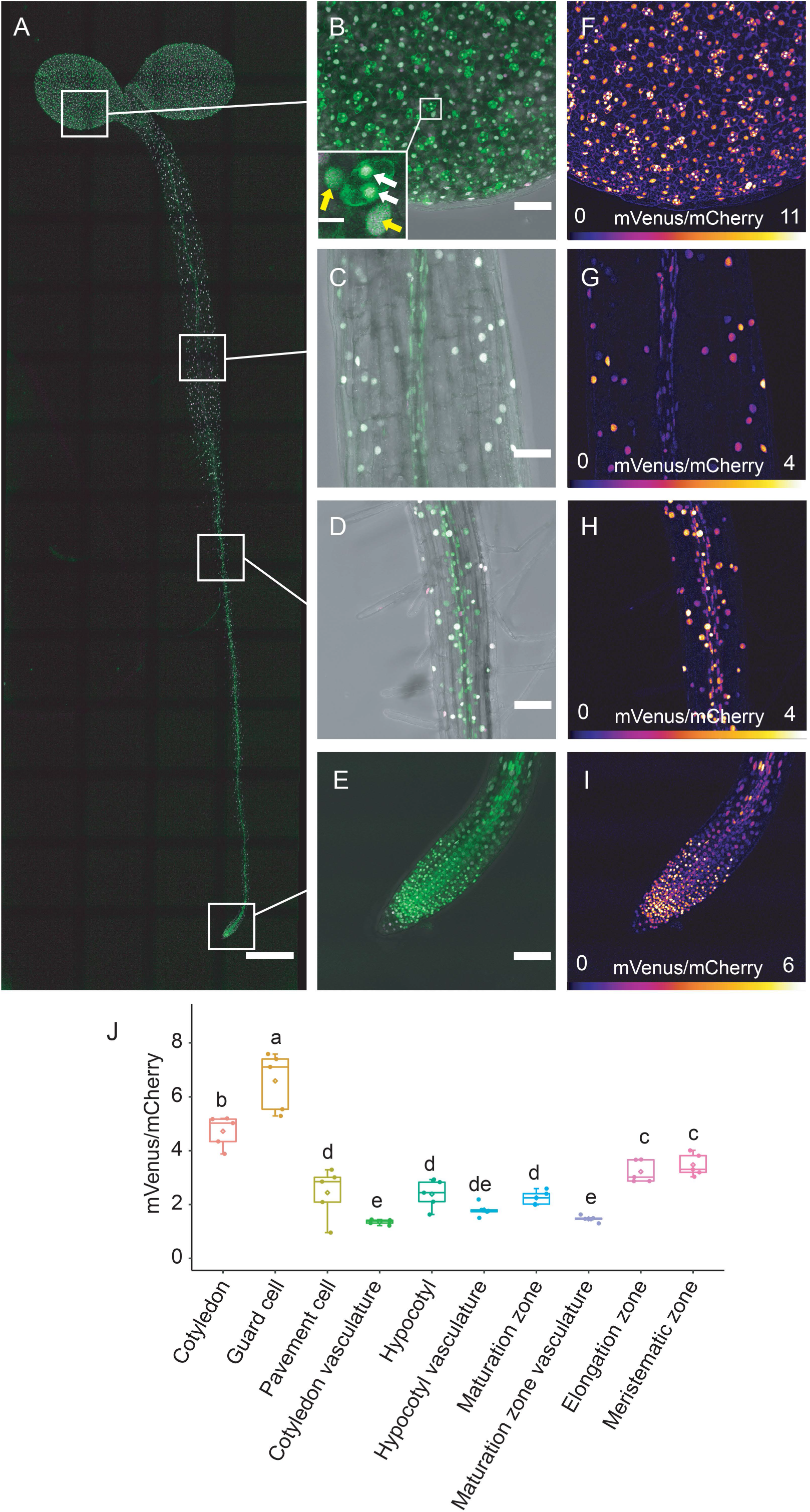
Expression of Strigo-D2 in Arabidopsis seedlings. **(A)** Ubiquitous activity of Strigo-D2. Shown are overlays of mVenus and mCherry-derived signals. Green: mVenus. Magenta: mCherry. Scale bar: 500 μm. **(B-E)** Activity of Strigo-D2 in cotyledons (B), hypocotyls (C), root maturation zones (D) and root tips (E). Images were recaptured from the positions indicated in (A) with higher magnification. Shown are overlays of mVenus, mCherry, and brightfield-derived signals. Green: mVenus. Magenta: mCherry. Grey: brightfield. Scale bars: 50 μm. White arrows in (B) indicate guard cells. Yellow arrows in (B) indicate pavement cells. Scale bar in the insert in (B): 10 μm. **(F-I)** Color-coded images generated from images shown in (B-E) visualizing mVenus/m-Cherry ratios indicating SL signaling levels. The color scale represents the range of intensity ratios among all targeted nuclei. 95th percentile of the ratios was used as the maximum value in order to eliminate outliers. **(J)** mVenus/m-Cherry signal ratios in different organs and tissues. Different letters above each box indicate statistical groups (One-way ANOVA, *p* < 0.01, n = 5).

### Strigo-D2 reveals tissue-specific (+)-5DS responsiveness

To challenge this conclusion, (+)-5DS was applied to cotyledons, hypocotyls, and the root maturation, elongation and meristematic zones. In cotyledons, a significant decrease of the mVenus/mCherry ratio was observed 20 min after application, and the ratio continued to decrease to a very low level within one hour (Fig 5A, Suppl. Movie 1). In comparison, ratio reduction was slower in hypocotyls where a significant decrease was only found 40 min after application (Fig 5B, Suppl. Movie 2). With 10 min after application, the fastest response was found in root maturation zones where a decrease of the ratio lasted until 70 min after the start of treatment (Fig 5C, Suppl. Movie 3). The sensor responded in a similar manner in root elongation zones, where a notable reduction of mVenus/mCherry ratios was observed at 20 min when ratios started to decrease sharply until reaching a minimum 40 min after application (Fig 5D, Movie 4). Interestingly, in the root apical meristem, a significant reduction of the mVenus/mCherry ratio was only found 40 min after (+)-5DS application (Fig 5E, Movie 4) arguing for a reduced responsiveness for this tissue. Indeed, when analyzing the root tip with higher spatial resolution along the longitudinal axis, we discovered a gradual decrease of responsiveness going from the maturation zone to the very tip of the root (Fig 6, A and B, Movie 5). Along the same lines, application of 0.5 and 0.05 µM (+)-5DS resulted in a similar Strigo-D2 response dynamics in the maturation and elongation zone, but the meristematic zone responded only slightly to 0.05 µM and not at all when adding 0.005 µM (+)-5DS (Fig 7A, B, and C). This was in contrast to the maturation and elongation zones which showed at least a slow response to 0.005 µm (+)-5DS (Fig 7A, B). These observations argued for a gradient of SL responsiveness in the root with a maximum in mature tissues and decreasing toward non-differentiated tissues along the longitudinal axis. Such a gradient is in line with a predominant expression of *D14* and *MAX2* in mature vascular tissues of the root (Fig S6) and the note that at least the D14 protein travels over short distances [31] potentially generating a concentration gradient of the capacity to perceive SL molecules.

**Fig 5.**
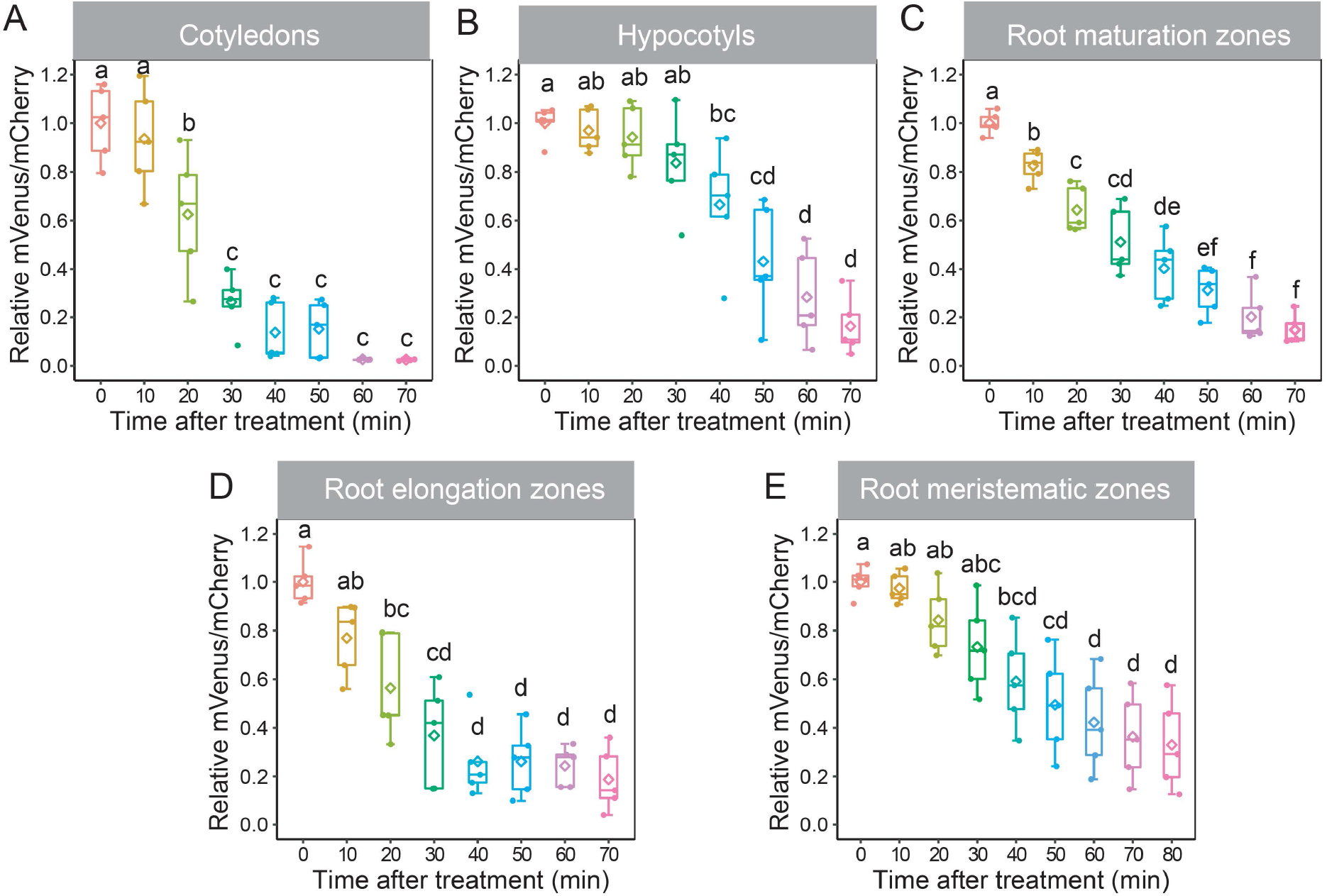
Strigo-D2 response to (+)-5DS treatment in different tissues. **(A)** Cotyledons. **(B)** Hypocotyls. **(C)** Root maturation zones. **(D)** Root elongation zones. **(E)** Root meristematic zones. Concentration of (+)-5DS: 0.5 μM. Age of the plant: 4 DAG. Different letters above each box indicate statistical groups (One-way ANOVA, *p* < 0.01, n = 5).

**Fig 6.**
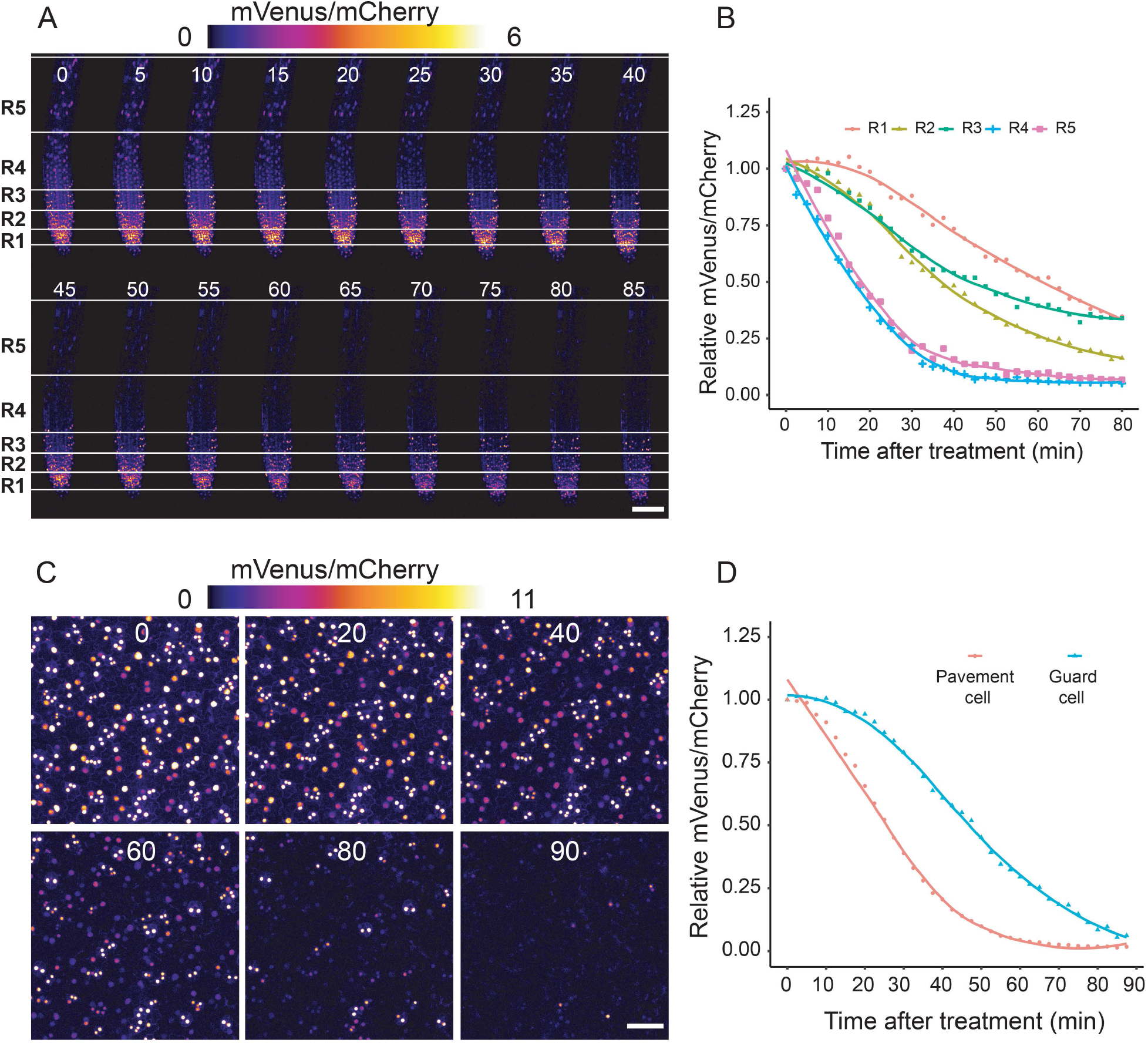
Strigo-D2 response to (+)-5DS at a cellular resolution in root tips and cotyledon epidermis. **(A)** Color-coded images of root tips generated from confocal images at the indicated time points after application. Concentration of (+)-5DS: 0.5 μM. Covered range of the regions (distance from bottom of meristematic zone): R1: 0-45 μm, R2: 45-90 μm, R3: 90-135 μm, R4: 135-435 μm, R5: 435-745 μm. Scale bar: 50 μm. **(B)** Dynamics of mVenus/mCherry ratios in four regions of the root tip after application. n = 3. **(C)** Color-coded images representing mVenus/mCherry signal ratios in the cotyledon epidermis at indicated time points after application. Concentration of (+)-5DS: 0.5 μM. Scale bar: 50 μm. **(D)** Dynamics of mVenus/mCherry ratios in pavement and guard cells. n = 3.

**Fig 7.**
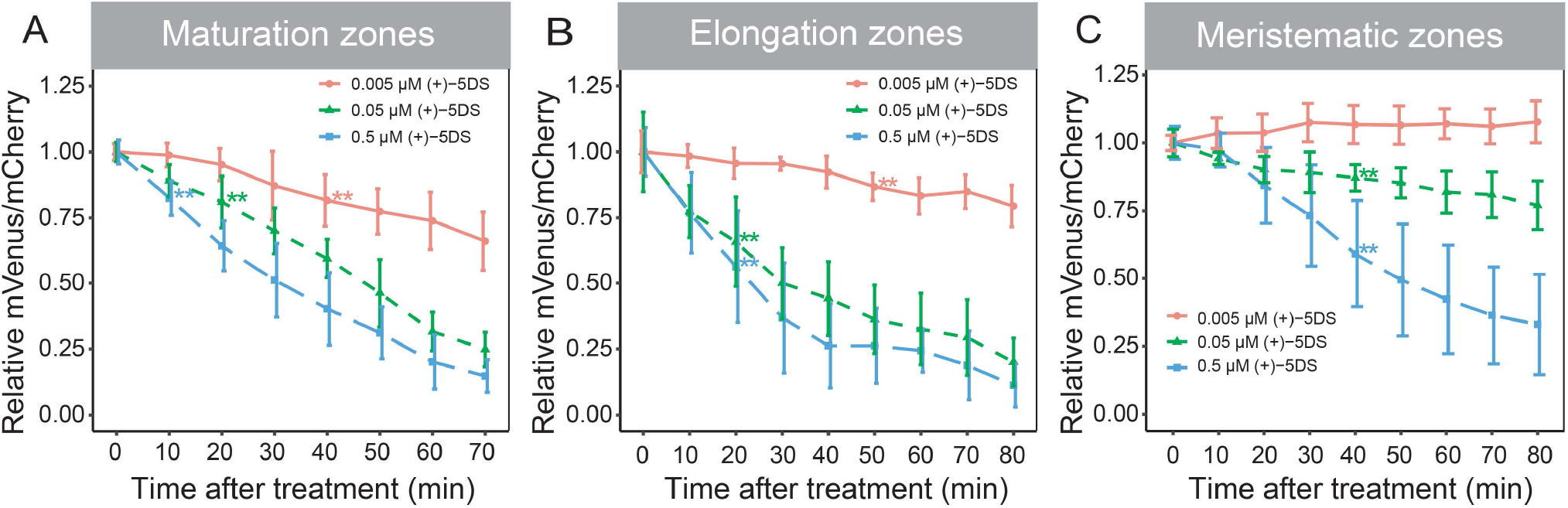
Sensitivity of Strigo-D2 to different (+)-5DS concentrations in root tips. **(A)** Maturation zones. **(B)** Elongation zones. **(C)** Meristematic zones. Asterisks (**) indicate the first time point of significant reduction of mVenus/mCherry ratios compared to before the treatment as determined by One-way ANOVA, *p* < 0.01. n = 5.

In cotyledons, pavement cells showed a rather fast response to (+)-5DS treatments and a sharp decrease of the mVenus/mCherry ratio. In comparison, the response in guard cells was relatively slow and lasted longer before the ratio reached a minimum level after 90 minutes (Fig 6C and 6D, Movie 6). This observation again argued for a cell type-specific SL signaling potential with substantial differences between neighboring cells. Interestingly, whereas transcriptional reporters revealed both *MAX2* and *D14* expression in the cotyledon vasculature, only *D14* expression could be detected in guard cells (Fig S6) implying that the a reduced Strigo-D2 response in these cells is particularly due to low levels of *MAX2* activity.

## Discussion

Genetically encoded biosensors are powerful tools for analyzing distribution and dynamics of small molecules with minimal invasion and high spatiotemporal resolution. Diverse biosensors for phytohormones or their activity have been successively developed over the past two decades (reviewed in [41]). However, ratiometric sensors allowing semi-quantitative analysis of strigolactone signaling in intact plants are still missing. As a major target of SL signaling in Arabidopsis, the full length SMXL6 protein has shown rapid *rac*-GR24-induced degradation in previous studies [47, 56], with some difficulties to detect a *35S*-driven SMXL6-YFP fusion in more differentiated tissues [57]. Similarly, in our study only weak mVenus signals were detected in transgenic *p35S:SMXL6-mVenus_p35S:mCherry-NLS* seedlings. Moreover, ubiquitous expression of a full SMXL6-mVenus protein had adverse effects on plant development making it unsuitable to serve as an informative readout for strigolactone signaling in a natural context. In comparison, the truncated SMXL6-D2-mVenus protein affected plant performance only slightly, was robustly detected in microscopic analyses and responded specifically to SL-signaling. In particular, D14-deficiency caused insensitivity of SMXL6-D2-mVenus against the pharmacological induction of SL-signaling whereas KAI2-deficiency had no effect. Although some cross-reaction of both pathways on the level of the SMXL protein targets has been suggested [35, 58], we thus conclude that SMXL6-D2-mVenus levels specifically report on SL signaling – as also supported by a large body of genetic and biochemical evidence for the SMXL6 protein in general [24, 47, 56].

Contradicting the previous conclusion that the D2 domain of a SMXL/D53 protein in rice is sufficient for protein interaction with D14 and for proteasomal degradation [22], exclusively the D1M domain of the Arabidopsis SMXL7 protein interacted with D14 in yeast-based and in transient split-LUC assays [35]. Together with the observations that D2 domains mediate homo- and heteromeric interactions between SMXL proteins and that a SMAX1-D2-LUC protein is not degraded in SMAX1/SMXL2-deficient backgrounds, these findings gave rise to the notion that isolated SMXL-D2 protein domains are only degraded due to their recruitment to SCF complexes by other full-length SMXL proteins [35]. If true, this model would predict that Strigo-D2 activity does not only depend on the presence of the SCF-complex components D14 and MAX2 but also on the availability of endogenous SMXL proteins interacting with the expressed SMXL6-D2-mVenus fusion protein. Because especially *SMXL6, SMXL7* and *SMXL8* promoter activities are, similar as promoter activities of *D14* and *MAX2*, associated with vascular tissues [24, 59, 60], this mode of action could contribute to the observed spatial pattern of Strigo-D2 activity with a minimum mVenus/mCherry ratio around the vasculature. However, because the formation of SMXL homo- and heteromers may, according to this concept, contribute to natural SL signaling, Strigo-D2 has the potential to report also on this aspect of the process. In addition, because isolated D2 domains interact with D14 at least *in vitro* in an SL-dependent fashion [22], a weak SL-triggered interaction of SMXL6-D2 with an SCF^MAX2/D14^ complex may already be sufficient to induce degradation without requiring interaction with other SMXL proteins. Moreover, as indicated by the severely altered phenotype of *p35S:SMXL6-mVenus_p35S:mCherry-NLS* plants, including the D1M domain into sensor construction may interfere with endogenous SL signaling events making the interpretation of sensor output difficult.

Independent from the mechanisms determining Strigo-D2 levels, we observed that, Strigo-D2 responds to exogenous SL analogs, without exception, in all cell types investigated. Considering the very local activity of some promoter reporters monitoring expression of genes encoding SL signaling components (this study, [29, 31]), this finding is remarkable as it suggests that, although to a different extent, all cells hold the potential for SL perception and signaling. With the caveat of the limitations of unraveling gene expression through promoter reporters, this means that expression of signaling components at low levels is sufficient or, as suggested previously for D14 [31], that SL signaling components travel at least short distances. Considering that also the expression of SMXL target proteins seems to be spatially rather restricted [24, 59], the question emerges whether a ubiquitous SL signaling capacity is indeed relevant and whether distinct SL responses are based on local or rather systemic effects. The recent identification of direct transcriptional targets of SMXL6, SMXL7 and SMXL8 proteins [39] allows analyzing target activity in respective mutants likewise with cellular resolution and, thereby, addressing this aspect. In fact, combination of the Strigo-D2 sensor with transcriptional reporters revealing target gene activity has the potential to reveal associations between signaling and signaling output. Establishment of a FRET-based direct sensor for SL molecules, as for example generated for gibberellins by employing the α/β-hydrolase-like receptor GIBBERELLIN INSENSITIVE DWARF 1A (GID1A) [61], will complete the toolbox for revealing the whole of the spatio-temporal complexity of the SL signaling process.

Importantly, different Strigo-D2 activities in different cell types argue for differences in the capacity of cells to respond to SLs. In our study, guard cells were on one end and vascular cells on the other end of the spectrum in this regard. In light of the reported role of SL signaling in increasing drought resistance [62-64] and the importance of long-distance transport of SL-related molecules possible along the vasculature [18], the relevance of these differences and their effect on distinct cell types is certainly interesting to investigate. Moreover, identifying factors limiting SL responsiveness in individual cells may provide means to modulate SL-dependent processes in a more targeted fashion. In guard cells, for example, we observed D14 but no MAX2 reporter activity prompting the question whether the absence of MAX2 causes a reduced response in this specific cell type. Considering its specificity and sensitivity toward SL signaling, the Strigo-D2 sensor may be useful for addressing this question. Another example for sensor utility may be the establishment of computational models for SL signaling patterns based on dynamic sensor outputs as done previously for gibberellin [65]. Overall, we assume that the dynamic nature of Strigo-D2 will facilitate quantitative determination of SL signaling for developing a deeper understanding of SL biology.

## Material and methods

### Plasmid construction

All constructs were generated via GreenGate cloning [66] if not mentioned otherwise. Used modules and primers are described in Supplemental Table S1 and Supplemental Table S2. Entry modules generated in this study were described as below. Three serial and partial fragments of the *SMXL6* coding sequence were amplified from cDNA with the primer pairs *SMXL6-1st-F* and *SMXL6-1st-R, SMXL6-2nd-F* and *SMXL6-2nd-R, SMXL6-3rd-F* and *SMXL6-3rd-R* to edit the endogenous Eco31I (BsaI) sites without altering the correspondent amino acid sequence. After digestion with Eco31I, the purified products were ligated into *pGGC000* [66]. We used the In-Fusion (Takara Bio) system to obtain the full-length *SMXL6* cDNA from these partial vectors with the following primer pairs (*E2-5-bp-deletion-F* and *E2-5-bp-deletion-R, SMXL6-frag1-into-E2-F* and *SMXL6-frag1-into-E2-R*). The *SMXL6-D2* fragment was cloned using primer pair *SMXL6-D2-F* and *SMXL6-3rd-R. D14* and *MAX2* promoters were amplified by primers described in Table S2 and cloned into *pGGA000* [66]. Entry modules with above mentioned desired fragments were subsequently cloned into intermediate modules using *pGGM000* by GreenGate reaction [66], and intermediate modules were finally transferred into destination vector *pGGZ000* [66, 67] via another GreenGate reaction.

### Plant material and generation of transgenic lines

In this study, the *Arabidopsis thaliana* Col-0 ecotype was used as genetic background. Loss-of-function alleles of *D14* (*d14-1*: WiscDsLoxHs137_07E) and *KAI2* (*htl-3*: 15 bp deletion [68]) were obtained from David Nelson (UC Riverside, US). Plants were transformed using an *Agrobacterium tumefaciens*-based floral-dip method to obtain transgenic lines.

### Plant growth condition

Seeds were surface-sterilized in a 1.5 ml microcentrifuge tubes with 700 μl of washing solution (70 % v/v ethanol in water, 0.1 % tween-20) by incubating in 37°C shaker (200 rpm) for 10 minutes, then washed 5 times with dH20 in the clean bench. Afterwards, seeds were stratified in 500 μl of dH_2_O at 4°C in the dark for 3 days and sown on plates with 0.8 % agar in 1/2 Murashige and Skoog (MS) medium supplemented with 1 % sucrose and grown vertically. Seedlings used for microscopic or phenotypic analysis were grown vertically in short-day conditions (SD, 10 h light and 14 h darkness) at 21°C. For microscopic analysis, seedlings were grown for four days after being transferred to growth chamber, while this period was increased to 5 days for root phenotypic analysis. For 4-week-old plants, from the third weeks onwards, seedlings were grown in long day (LD, 16 h light and 8 h dark) conditions at 21°C. For the independent transgenic lines in WT background, only lines with single integration based on T2 segregation ratios were propagated to T3, and lines homozygous for the Basta resistance were selected for characterization.

### Chemicals and Treatments

The SL substrates (+)-5-deoxy-strigol [(+)-5DS, CAS No: 151716-18-6] was obtained from OlChemIm Ltd (Olomouc, Czech Republic), (±)-GR24 (*rac*-GR24, CAS No: 76974-79-3) and Karrikin1 (KAR1, CAS No: 857054-02-5) were purchased from Chiralix (Nijmegen, Netherland). GR24^4DO^ was obtained from StrigoLab (Torino, Italy). 3-Indoleacetic acid (IAA, CAS No: 87-51-4), trans-Zeatin (cytokinin, CK, CAS No: 1637-39-4), (+/-)-Abscisic acid (ABA, CAS No:14375-45-2), and Gibberellic acid (GA_3_, CAS No: 77-06-5) were purchased from Merck KGaA/ Sigma-Aldrich (Darmstadt, Germany). For generating stock solutions, (+)-5DS, *rac*-GR24, KAR1, and GR24^4DO^ were dissolved in acetone, IAA, CK, and ABA were dissolved in 1 N NaOH and GA_3_ was dissolved in ethanol before diluted with water to reach stock concentrations. All hormone and mock solutions were then diluted with 1/2 MS medium to working concentrations. The same working concentration of 0.5 μM was applied for (+)-5DS, *rac*-GR24, KAR1, and GR24^4DO^ if not stated otherwise. Working concentrations for IAA, CK, ABA, and GA_3_ were 1, 5, 2, and 5 μM, respectively. Five 4-day-old seedlings were transferred from 1/2 MS plates onto a chambered coverslip (Cat.No:80286, ibidi GmbH, Gräfelfing, Germany) containing 120 μl treatment solution. The seedlings were immersed in solution and covered by a piece of fabric mesh. Then the slide chamber was covered with lid to avoid evaporation.

### Preparation for time-lapse imaging

Four-day-old seedling were transferred onto a chambered coverslip containing 60 μl 1/2 MS medium, covered by a piece of mesh, and fixed by an adjusted paper clip to avoid movement. The slide chamber was covered by a glass slide and sealed with water to avoid evaporation. Afterwards, the position of interest on the seedling was marked under the microscope and an image for time point zero (t0) was acquired. Then, 60 μl of (+)-5DS was added through the mesh and mixed with the 1/2 MS medium by gently pipetting and recording was started immediately. Focus was adjusted before the first timepoint of time-lapse imaging t1 and the time-lapse imaging was started from t1. Effect of different concentrations was monitored in root maturation zones, elongation zones, and meristematic zones using final concentrations of 0.5, 0.05, and 0.005 μM (+)-5DS.

### Image acquisition and analysis

Bright field and fluorescence images were acquired using a Leica TCS SP8 confocal microscope (Leica Microsystems, Wetzlar, Germany), mVenus was excited by 514 nm laser light, while mCherry was excited using 561 nm. Emissions were detected sequentially to avoid cross-talk between fluorophores. mVenus and mCherry fluorescence were detected in the range of 524-540 nm and 571-630 nm, respectively. The cyan fluorescent protein (CFP) in *D14* and *MAX2* reporter lines was excited by 458 nm laser light and detected in the range of 465-509 nm. The yellow fluorescent protein (YFP) in the SMAX1-YFP reporter line was excited by 514 nm laser light and detected in the range of 524-540 nm. The output power of the 514 nm laser was set to 30 %. For imaging of seedlings expressing SMXL6-D2-mVenus and mCherry-NLS, the intensity of 514 nm and 561 nm lasers were consistently set to 1.0 % and 0.04 %, respectively. While for imaging of seedlings expressing SMXL6-mVenus and mCherry-NLS, the intensity of 514 nm and 561 nm lasers were consistently set at 10 % and 0.04 %, respectively. Tile scanning of whole seedlings was performed using a 10x objective, while other images were acquired using a HC PL APO 20x/0,75 IMM objective. Z-stacks covering the entire thickness of observed tissue were acquired. Step size for tile scanning was 10 μm and that for other image acquisitions was 5 μm. Images were analyzed using Fiji [69]. Maturation, elongation and meristematic zones of roots were defined as previously described [46]. For determining root length, plates were scanned by an Epson Perfection V600 Photo scanner and measured in Fiji using the freehand line tool. Composite images containing mVenus and mCherry channel information were generated by Fiji and automatically processed and analyzed using macro codes. For image stacks, Z projections using the “maximum projection” option were performed. Composite images with multiple frames (generated from time-lapse imaging) were saved as image sequence before intensity measurements. To measure fluorescence intensity in each individual channel, proper intensity thresholds were set depending on the targeted cell type to eliminate noise from auto-fluorescence. The “Watershed” function was applied to separate crowded nuclei. Afterwards, nuclei were sampled and measured using the “Analyze Particles” tool for each time point and each channel separately. For images containing different types of cells, the “freehand” tool was used to select regions of specific cells. Fluorescence intensity of each nucleus was determined using gray values collected in the respective channels. Fluorescence intensity ratio of each nucleus was calculated in Excel (Microsoft, Redmont, US) by dividing the values of mVenus by the values of mCherry. The mean value of all nuclei in the same frame was taken as one biological repetition. Relative fluorescence intensity ratio references the average ratio at time 0 min or to the mock, respectively. The same method was applied for all images of each analysis. False color images were generated by “Calculator Plus” in Fiji through calculating intensity ratios of each pixel from mVenus and mCherry channels after being Gaussian Blurred and subtracting background. Color scale was calculated based on the range of intensity ratios of all the nuclei. The 95^th^ percentile of the ratios was used as the maximum value in order to eliminate outliers. One-way analysis of variance (ANOVA) was conducted using R, statistical significance among data sets was calculated using the Duncan’s test. Charts were generated in R using ggplot2 [70].

## Supporting information

### Supplementary figures

**Fig S1**. Comparison of fluorescent signals in transgenic seedlings carrying Strigo-D2 or *p35S:SMXL6-mVenus_p35S:mCherry-NLSs* transgenes.

**Fig S2**. Effect of SLs and KAR1 on *pSMXL5:SMAX1-YFP* reporter activity.

**Fig S3**. Strigo-D2 response to (+)-5DS and other plant hormones in the root maturation zone.

**Fig S4**. Strigo-D2 activity in seedlings.

**Fig S5**. Strigo-D2 activity in the vasculature of cotyledons, hypocotyls and root maturation zones.

**Fig S6**. *D14* and *MAX2* promoter reporter activity in seedlings.

### Supplementary Tables

**Table S1:** GreenGate Vectors used in this study

**Table S2:** Primers used in this study

### Supplementary movies

**Movie 1: Strigo-D2 response to (+)-5DS treatment in cotyledons**. Shown are overlays of mVenus and mCherry-derived signals. Green: mVenus. Magenta: mCherry. Time after (+)-5DS treatment is indicated at the upper left corner of the movie. Scale bar: 50 μm. Same settings were applied in Movies 2, 3, 4, and 6.

**Movie 2: Strigo-D2 response to (+)-5DS treatment in hypocotyls**.

**Movie 3: Strigo-D2 response to (+)-5DS treatment in root maturation zones.**

**Movie 4: Strigo-D2 response to (+)-5DS treatment in root tips**.

**Movie 5: Gradual decrease of Strigo-D2 responsiveness to (+)-5DS from the maturation zone to the very tip of the root**. Shown are overlays of mVenus, mCherry, and brightfield-derived signals. Green: mVenus. Magenta: mCherry. Grey: brightfield. Time after (+)-5DS treatment is indicated at the upper left corner of the movie. Scale bar: 50 μm.

**Movie 6: Comparison between Strigo-D2 response to (+)-5DS in pavement and in guard cells**.

## Acknowledgments

This work was supported by a fellowship of the Chinese Scholarship Council (CSC) and a Peterson scholarship (http://www.petersonelites.sdu.edu.cn/jxjjj.htm) to J.Z., an ERC Consolidator grant [647148, PLANTSTEMS] and a grant of the Deutsche Forschungsgemeinschaft [DFG, GR_2104/4-1] to T.G., postdoctoral fellowships of the Alexander von Humboldt-Stiftung [3.5-JPN –1164674-HFST-P] and the Japan Society for the Promotion of Science [JSPS Overseas Research Fellowships 201960008] to D.S.. We thank the Nikon Imaging Center (NIC) of Heidelberg University for support.

## Author Contributions

Designed and conducted experiments CSo, JZ, MG, DS, CSc, VJ. Conceptualized experiments: GG, TG. Wrote the manuscript: CSo, TG.

