## Supplementary material for "Strigo-D2 – a bio-sensor for monitoring the spatio-temporal pattern of strigolactone signaling in intact plants": Fig S2

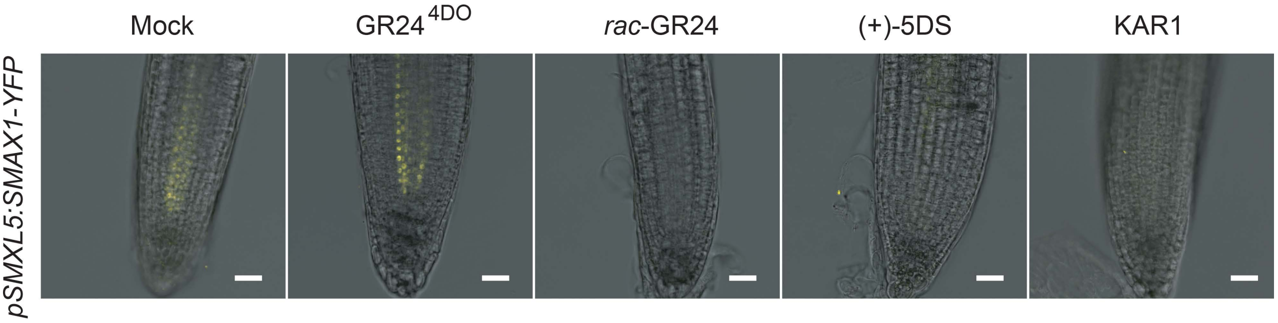


**Fig S2. Effect of SLs and KAR1 on pSMXL5:SMAX1-YFP reporter activity.** YFP-derived signal visualized in yellow. Treatment duration: 30 min. Scale bars: 50 µm.
