## Supplementary material for "Strigo-D2 – a bio-sensor for monitoring the spatio-temporal pattern of strigolactone signaling in intact plants": Fig S3

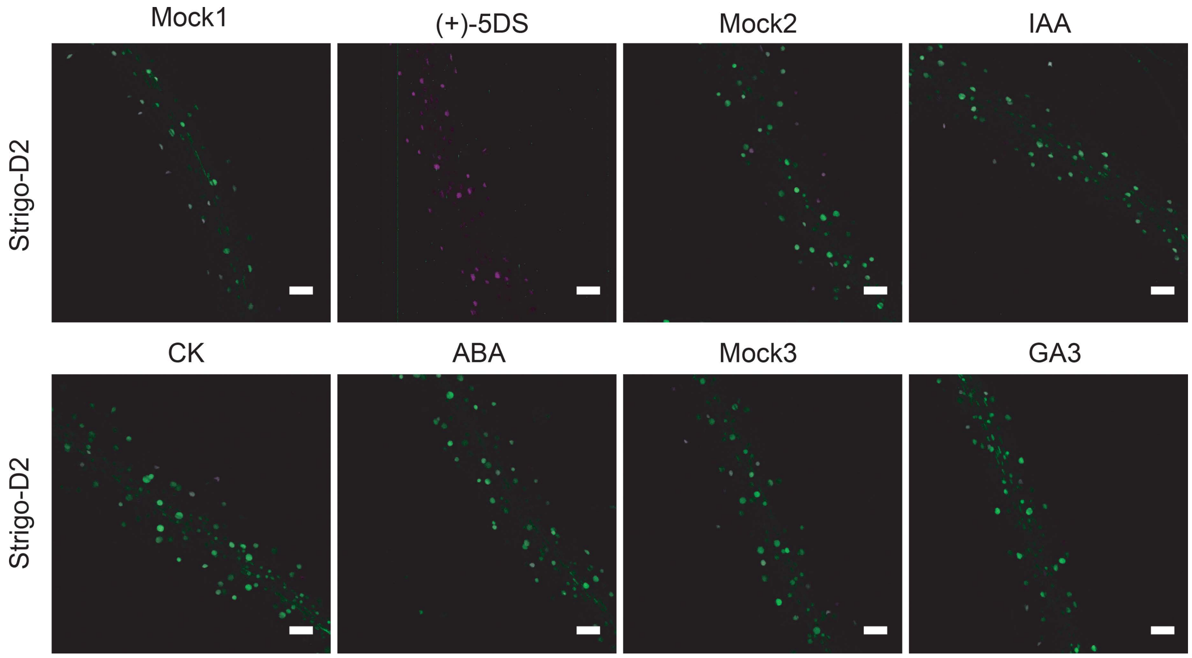


**Fig S3. Strigo-D2 response to (+)-5DS and other plant hormones in the root maturation zone.** Images merging mCherry (magenta) and mVenus (green)-derived signals are shown. Treatment duration: 120 min. Mock1: treatment with (+)-5DS solvent. Mock2: treatment with IAA, trans-Zeatin, and ABA solvent. Mock3: treatment with GA3 solvent. Scale bars: 50 μm.
