## Supplementary material for "Strigo-D2 – a bio-sensor for monitoring the spatio-temporal pattern of strigolactone signaling in intact plants": Fig S4

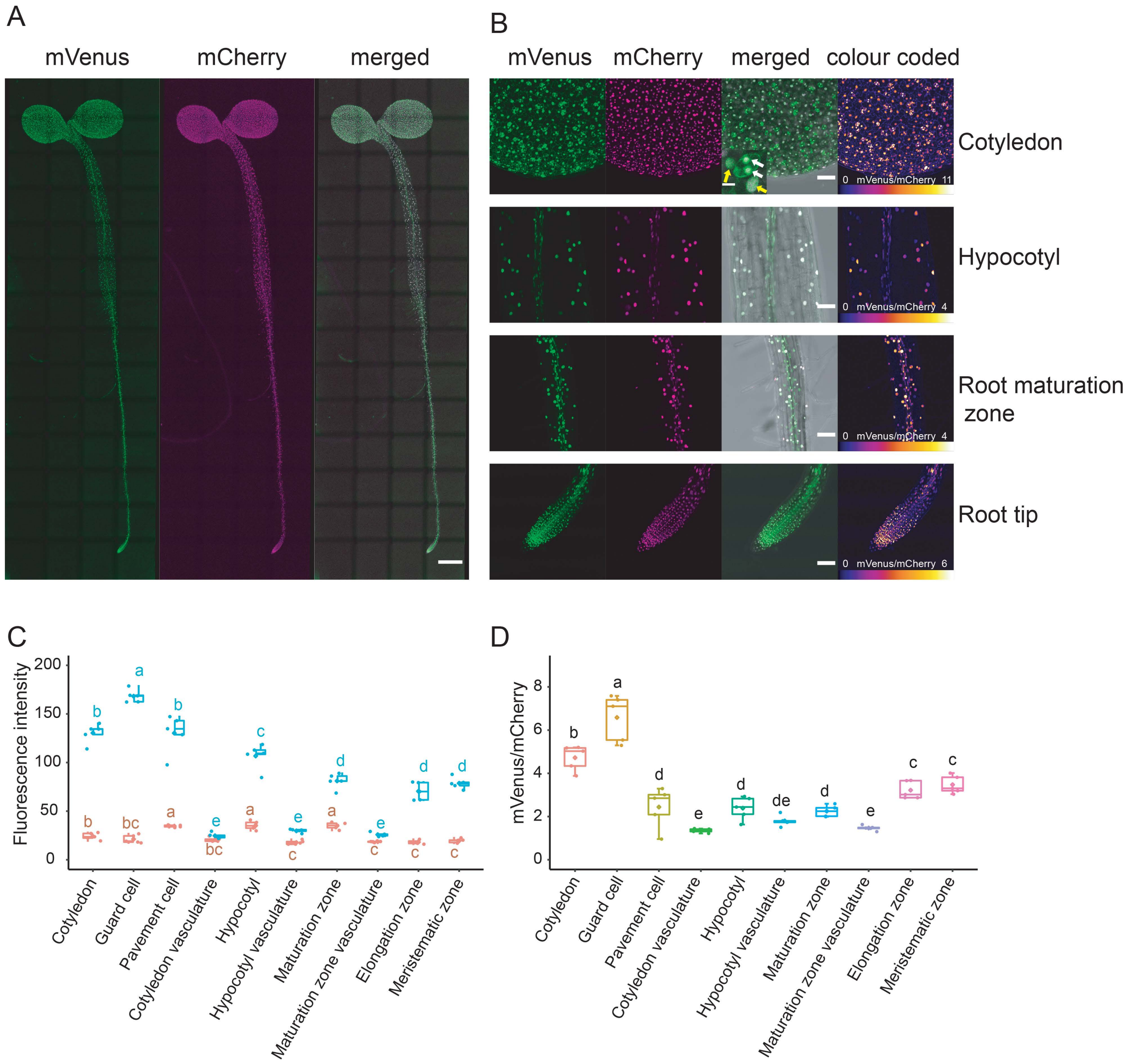


**Fig S4. Strigo-D2 activity in seedlings. (A)** Ubiquitous expression of Strigo-D2 in seedlings. Scale bar: 500 μm. **(B)** Activity of Strigo-D2 in cotyledons, hypocotyls, root maturation zones and root tips. Images were recaptured from corresponding positions in (A) with higher magnification. Scale bars: 50 μm. White arrows: guard cells. Yellow arrows: pavement cells. Scale bar in magnified image: 10 μm. **(C)** Fluorescence intensity of mVenus and mCherry detected in different tissues or cells. **(D)** Intensity ratios of mVenus and mCherry in different tissues or cells.
