## Supplementary material for "Strigo-D2 – a bio-sensor for monitoring the spatio-temporal pattern of strigolactone signaling in intact plants": Fig S5

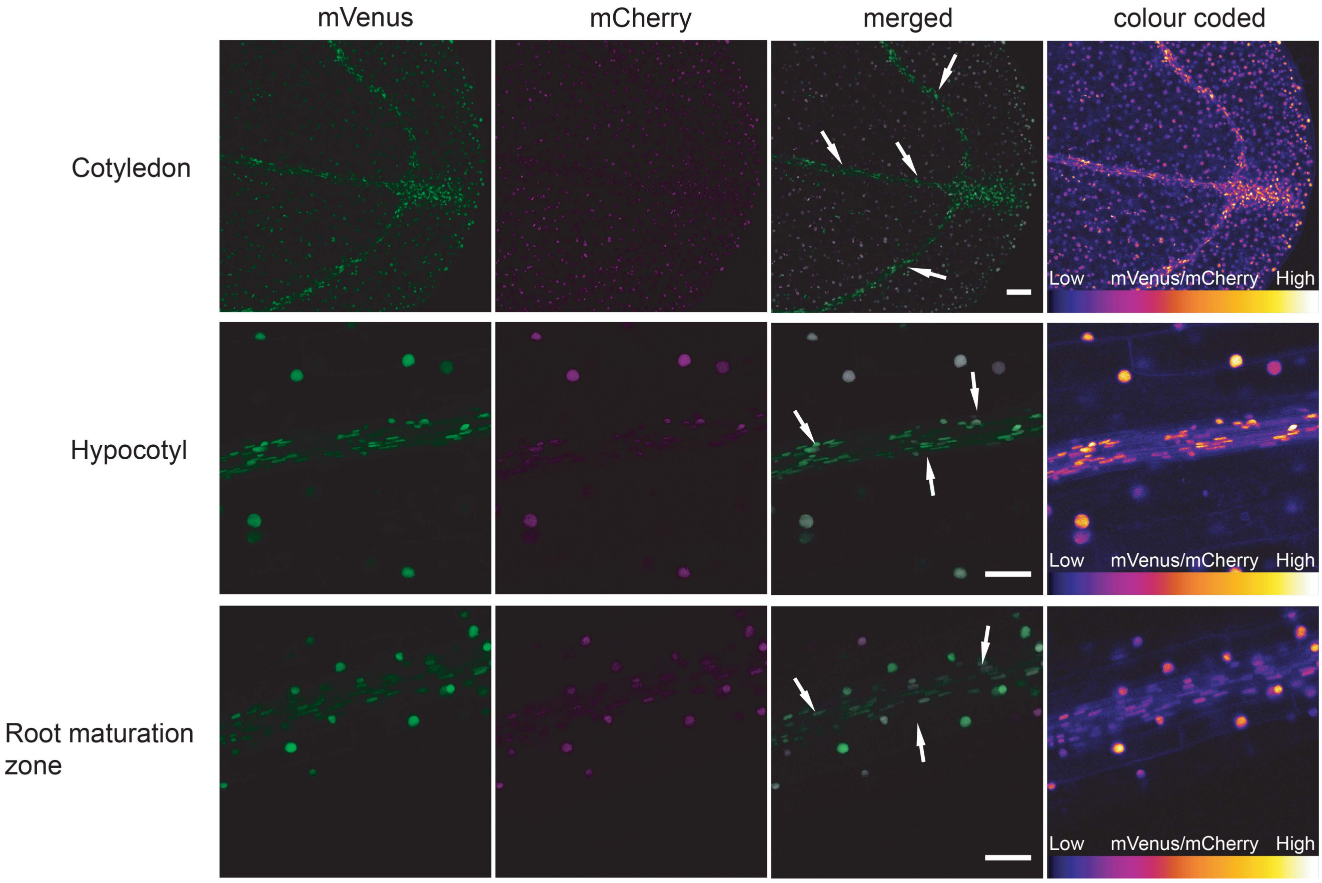


**Fig S5. Strigo-D2 activity in the vasculature of cotyledons, hypocotyls and root maturation zones.** White arrows indicate vascular strands. Scale bars: 50 µm.
