## Supplementary material for "Strigo-D2 – a bio-sensor for monitoring the spatio-temporal pattern of strigolactone signaling in intact plants": Fig S6

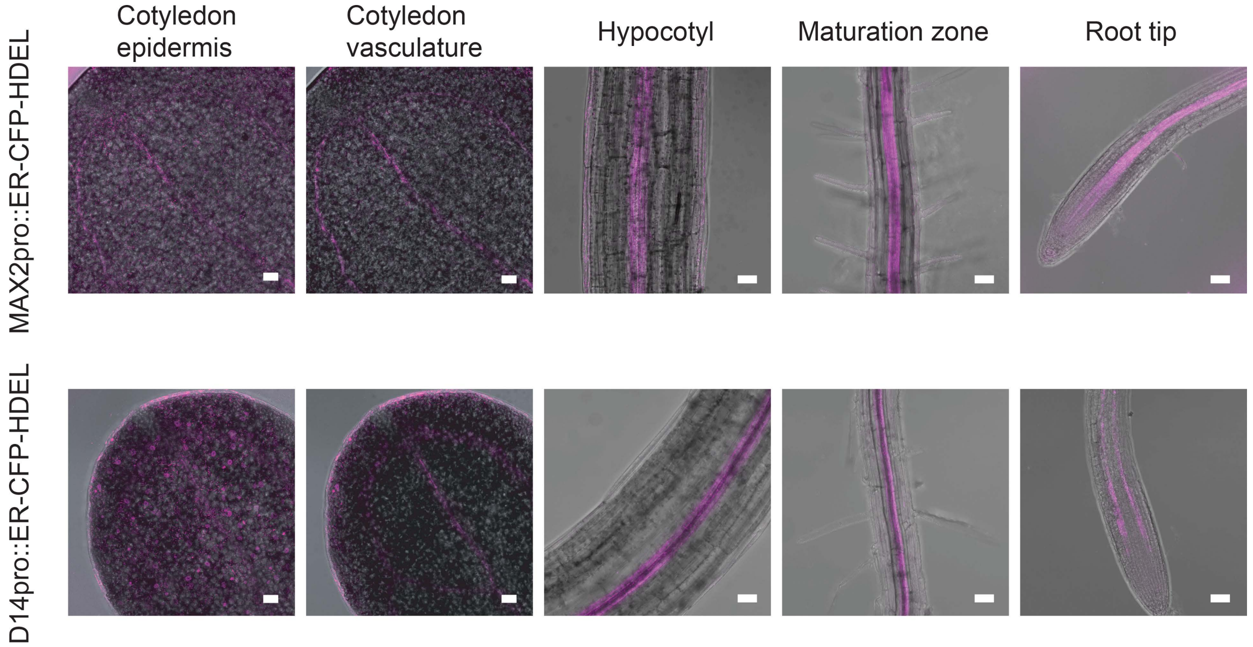


**Fig S6. *D14* and *MAX2* promoter reporter activity in seedlings.** CFP fluorescence is visualized in magenta. Scale bar: 50 μm.
