## Supplementary material for "Strigo-D2 – a bio-sensor for monitoring the spatio-temporal pattern of strigolactone signaling in intact plants": Table S1

**Table S1. GreenGate Vectors used in this study**

| Vector name | Vector ID | Modules |
|---|---|---|
| *p35S:SMXL6-mVenus_p35S:mCherry-NLS* | *pCS12* | *35S* (*pGGA004**), B-dummy (*pGGA022**), *pGGC-SMXL6* (*pCS1*), *pGGD-mVENUS* (*pD00587* ‡), *rbscT* (*pGGE001**), F-H adapter (*pGGG001**), H-A adapter (*pGGG002**), *mCherry* (*pGGC015**), *NLS* (*pGGD007**), *BastaR* (*pGGF001**), Destination vector (*pGGZ003**) |
| *p35S:SMXL6-D2-mVenus_p35S:mCherry-NLS* | *pJZ27* | *35S* (*pGGA004**), B-dummy (*pGGA022**), *pGGC-SMXL6-D2* (*pJZ25*), *pGGD-mVENUS* (*pD00587* ‡), *rbscT* (*pGGE001**), F-H adapter (*pGGG001**), H-A adapter (*pGGG002**), *mCherry* (*pGGC015**), NLS (*pGGD007**), *BastaR* (*pGGF001**), Destination vector (*pGGZ003**) |
| *D14pro:ER-CFP_WOXpro:ER-YFP* | *pVJ33* | *D14pro* (*pVJ25*), ER Signal Peptide (*pGGB006**), *mTurquoise2* (*pSW596*†), HDEL (*pGGD008**), *ter AT4G24550* (*pVL12*†), F-H adapter (*pGGG001**), H-A adapter (*pGGG002**), *WOX4pro* (*pVL37*†), *mVenus* (*pSW549*†), *tWOX4* (*pVL22*†), *BastaR* (*pGGF001**), Destination vector (*pGGZ003**) |
| *MAX2pro:ER-CFP_ WOXpro:ER-YFP* | *pVJ47* | *MAX2pro* (*pVJ40*), ER Signal Peptide (*pGGB006**), *mTurquoise2* (*pSW596*†), HDEL (*pGGD008**), *ter AT4G24550* (*pVL12*†), F-H adapter (*pGGG001**), H-A adapter (*pGGG002**), *WOX4pro* (*pVL37*†), mVenus (*pSW549*†), *tWOX4* (*pVL22*†), *BastaR*, (*pGGF001**), Destination vector (*pGGZ003**) |

*reference: [66]

†reference: [67]

‡reference: [71]
