## Supplementary material for "Strigo-D2 – a bio-sensor for monitoring the spatio-temporal pattern of strigolactone signaling in intact plants": Table S2

**Table S2. Primers used in this study**

| Primer name | Sequence (5’-3’) | Usage |
|---|---|---|
| *SMXL6-1st-F* | AACAGGTCTCAGGCTATGCCGACGCCGGTGACTACGG | Cloning 1st part of *SMXL6* into *pGGC000* |
| *SMXL6-1st-R* | AACAGGTCTCTCGCCTGTGACCGTTTGATCGCCGC | Cloning 1st part of *SMXL6* into *pGGC000* |
| *SMXL6-2nd-F* | AAACGGTCTCAGGCGAACCAGAG | Cloning 2nd part of *SMXL6* into *pGGC000* |
| *SMXL6-2nd-R* | TTCTTTGGTCTCAATAGCCCGTAGCC | Cloning 2nd part of *SMXL6* into *pGGC000* |
| *SMXL6-3rd-F* | AACAGGTCTCGCTATTGAAACCAAAGAAGACAAGGGAATAACAGGC | Cloning 3rd part of *SMXL6* into *pGGC000* |
| *SMXL6-3rd-R* | AACAGGTCTCACTGACCATATCACATCCACCTTCGCC | Cloning 3rd part of *SMXL6* into *pGGC000* and *SMXL6-D2* cloning |
| *E2-5-bp-deletion-F* | GCTTTAGATGACGCTAATACATCAGCC | In fusion of *SMXL6* parts |
| *E2-5-bp-deletion-R* | CTTACTGCTGCCTGTTATTCCCTTG | In fusion of *SMXL6* parts |
| *SMXL6-frag1-into-E2-F* | TGAAGCTTGGTCTCAGGCTATGCC | In fusion of *SMXL6* parts |
| *SMXL6-frag1-into-E2-R* | CGTCTCTGGTTCGCCTGTGACCGTTTGATCGCCG | In fusion of *SMXL6* parts |
| *SMXL6-D2-F* | AACAGGTCTCAGGCTCAATGCAGAAAGATTTCAAGTCTC | *SMXL6-D2* cloning |
| *pMAX2_ModuleA F* | AACAGGTCTCAACCTTGGAACGAACGTGGAGATCG | *MAX2* promoter cloning |
| *pMAX2_ModuleA R* | AACAGGTCTCATGTTGAGAAGCGGCAAATCTACAA | *MAX2* promoter cloning |
| *pD14_ModuleA F* | AACAGGTCTCAACCTAAATGTCTTAACCATCTTAA | *D14* promoter cloning |
| *pD14_ModuleA R* | AACAGGTCTCATGTTTTTTTTATGTGTTTGGGTTTG | *D14* promoter cloning |
