## Supplementary material for "Strigo-D2 – a bio-sensor for monitoring the spatio-temporal pattern of strigolactone signaling in intact plants": Fig S1

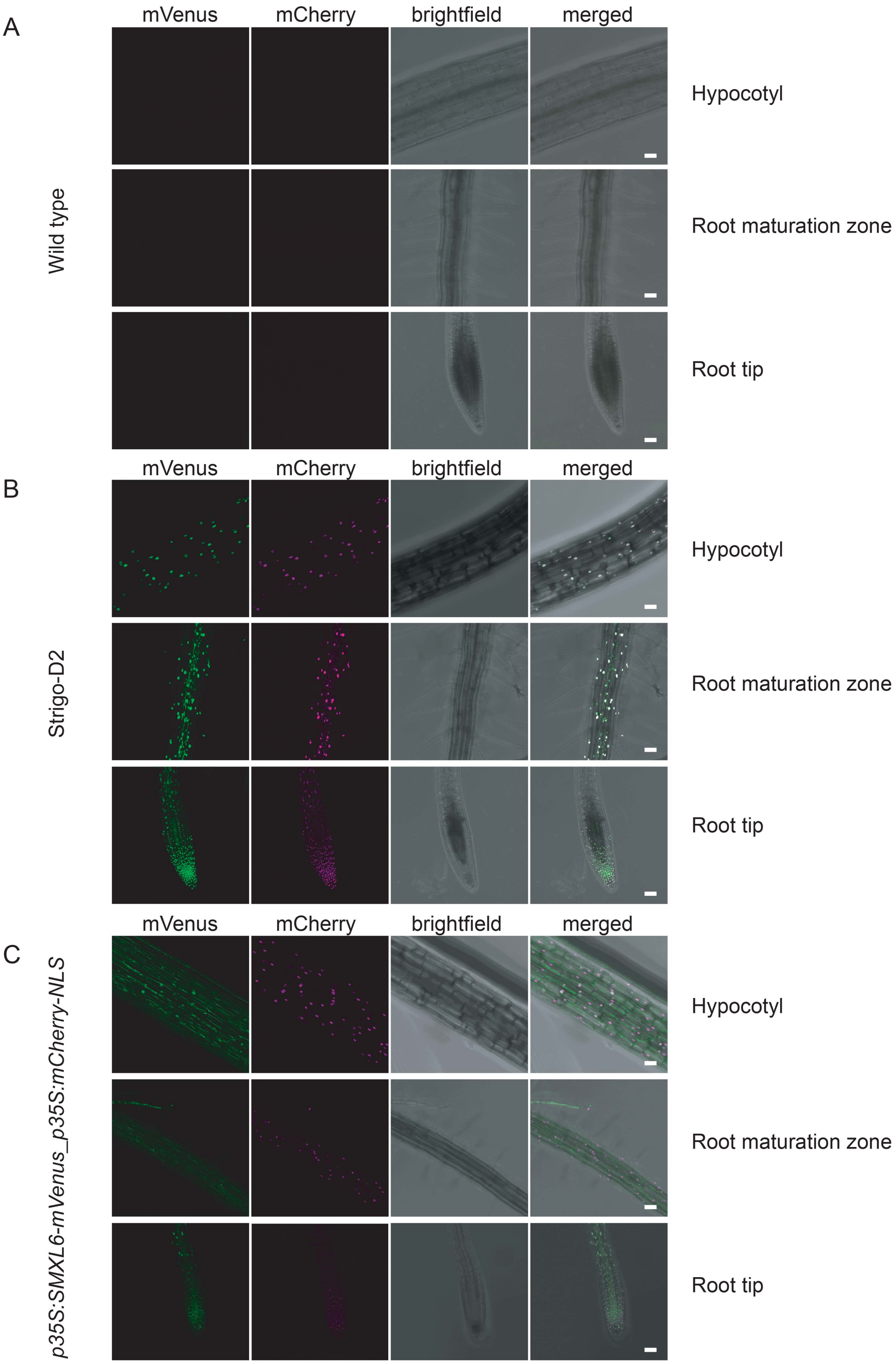


**Fig S1. Comparison of fluorescent signals in transgenic seedlings carrying Strigo-D2 or p35S:SMXL6-mVenus_p35S:mCherry-NLS transgenes. (A)** Wild type seedling without transgene. Laser excitation intensity for mVenus: 1 %, Detection wavelength range 519-555 nm. **(B)** Strigo-D2 seedling. Laser excitation intensity for mVenus: 1 %, Detection wavelength range 524-540. **(C)** *p35S:SMXL6-mVenus_p35S:m-Cherry-NLS* seedling. Laser excitation intensity for mVenus: 10 %, Detection wavelength range 519-555. Scale bars: 50 μm.
